# Modeling framework disentangles cerebellar mechanisms in speech feedback control, revealing trade-offs reshaped by degeneration

**DOI:** 10.64898/2026.07.14.738521

**Authors:** Alvincé L Pongos, Kwang S. Kim, Jessica L. Gaines, Vikram Ramanarayanan, Nefeli Chanoutsi, Rabab Rangwala, Kurtis Brent, Benjamin Parrell, John F. Houde, Srikantan S. Nagarajan

## Abstract

A central challenge in systems neuroscience is understanding how computational mechanisms—including those implemented within a single brain region—interact to produce behavior. For example, prior work in the literature attributes many computational functions to the cerebellum, but these functions have been tested in isolation and it remains unclear how they jointly contribute to motor control. Here, we test several established hypotheses of cerebellar function: internal modeling, timing of movement dynamics, sensory-error processing, delay processing, and multimodal integration. We first formalize these functions as mechanistic parameters within a computational model of speech motor control. We then use this formalism to investigate the relative contribution of each function to the abnormal speech corrective response seen in adults with cerebellar degeneration during perturbed auditory feedback. We find the following functions explain most of the behavioral differences: internal modeling, timing of movement dynamics, and multimodal integration. We also show that the key mechanisms have a trade-off relationship, and that cerebellar degeneration modulates those trade-off strengths and boundaries. These results both elaborate the mechanistic function of the cerebellum in speech feedback control and, more broadly, demonstrate the promise of using this paradigm to simultaneously test competing theories of neural function underlying behavior.

## Introduction

A central goal of systems neuroscience is to understand how computations across neural networks–including computations within a single brain region–interact to generate behavior. For example, the neural control of reaching, walking, or speaking depends on coordinated computations across cortical, subcortical, and cerebellar circuits [1, 2, 3, 4]. Within this brain network, high-level computations include predicting the consequences of actions [5, 6], accounting for delays and noise in sensory feedback [7, 8], and correcting ongoing movements when predicted and observed feedback diverge [9, 10, 11]. Within this network, the cerebellum itself is thought to support several distinct computations, including internal modeling [9, 7, 8, 12, 13], sensory-error processing [14, 15, 16], multimodal sensory integration [17, 18, 19, 20, 21], delay processing [22, 23, 24], and the timing of movement dynamics [25, 26, 27, 28]. However, because these computations are generally examined in isolation and can produce similar corrective behavior, it remains difficult to determine how they operate together during motor control. To help address this gap, we use speech production as a model system, focus on cerebellar computations, and formalize these computations as mechanistic parameters in a model of speech motor control for the following reasons. The first reason is that the cerebellum is known to causally contribute to speech motor behavior [29, 30, 31, 32]. The second reason is that computational models of speech motor control can reproduce speech behavior with high fidelity [33, 34, 35]. The third reason is that although cerebellar hypothesized functions have been well studied [9, 15, 24, 27], the relative contributions of candidate cerebellar computations to disordered speech remain unresolved. Therefore, in this work we formalize cerebellar computations under a single computational framework and investigate: which cerebellar computational mechanisms contribute to corrective speech behavior, what their relative contributions are, and how they interact to produce abnormal corrections observed in adults with cerebellar degeneration.

Multiple lines of evidence—lesion, stimulation, and neurodegenerative studies—show that the cerebellum plays a critical role in healthy speech production, although the precise functions of the cerebellum remain unclear. Cerebellar damage often leads to speech motor impairments: early cerebellar lesion studies, including Holmes (1917) and Zentay (1937)[29, 30], report symptoms such as slow, monotonous, jerky, slurred, and labored speech. Later work, including individuals with both focal lesions and diffuse cerebellar degeneration, further systematized these observations [36, 37]. They showed that ataxic dysarthria (the speech motor disorder associated with cerebellar damage), regardless of the precise etiology, is characterized by prosodic, timing, and coordination disruptions, along with imprecise spatial control and, more variably, vocal and respiratory irregularities. One potential explanation for at least a subset of these impairments is that impaired predictive, feedforward models lead to an increased reliance on reactive, feedback control [38, 39]. Consistent with this hypothesis, several studies show that individuals with cerebellar degeneration (CD) produce larger compensatory speech responses to external perturbations of auditory feedback during speaking, compared to healthy controls (HCs) [38, 40, 41], though these differences are not always observed (Parrell et al., 2021 [42]). Consistent with these results, non-invasive neurostimulation of the cerebellum in healthy participants alters compensation to external perturbations of vocal pitch perturbations [31, 32], though again these effects appear protocol-dependent and are not always observed [43]. Taken together, these studies indicate cerebellar damage may increase the reliance on feedback control and, as a consequence, increase the magnitude of the corrective, feedback-driven responses to externally introduced sensory errors. However, this behavior could arise through changes to several of the hypothesized neurocomputational functions (Table 1), making the precise mechanisms driving these changes unclear.

**Table 1.**
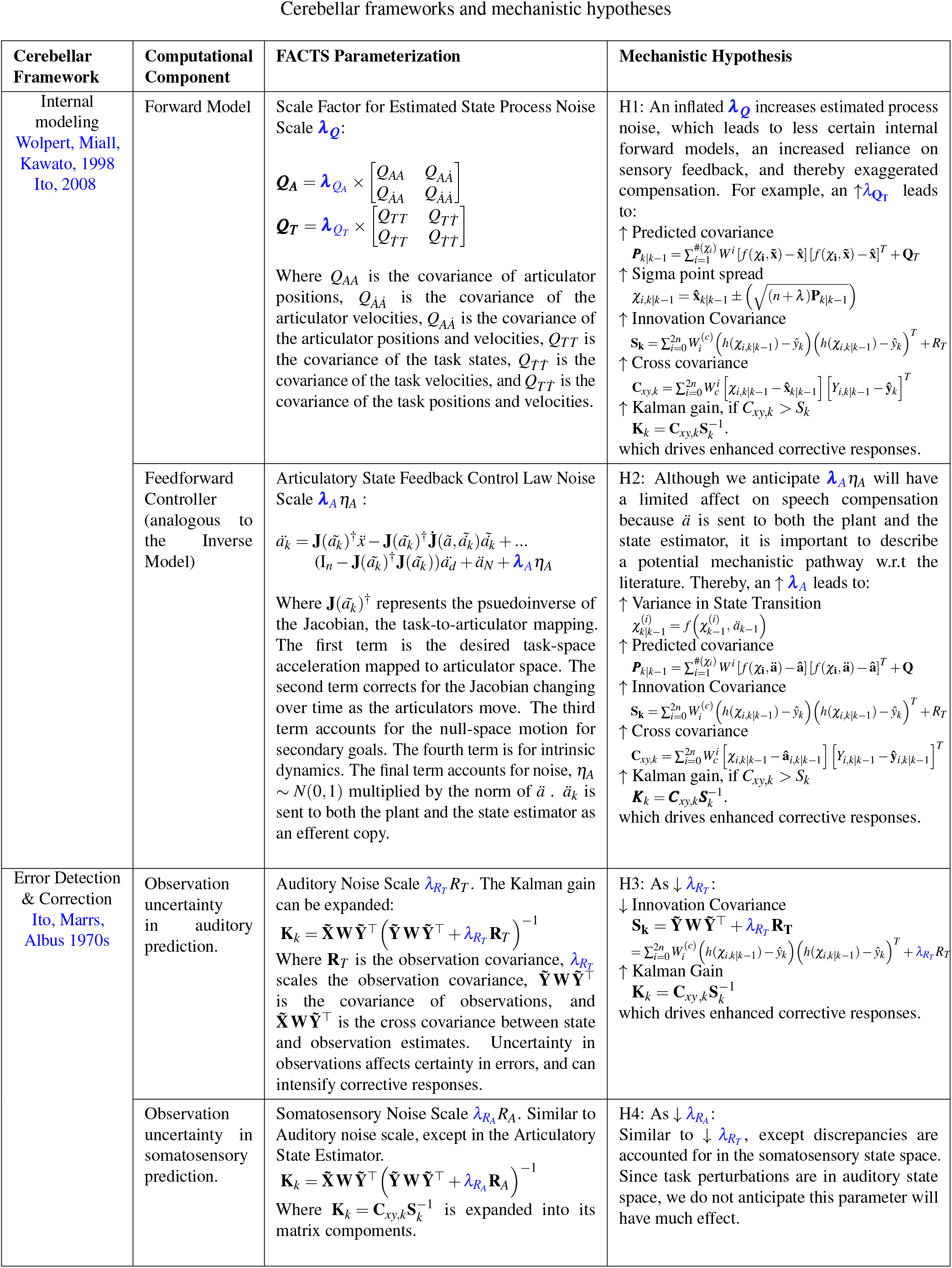
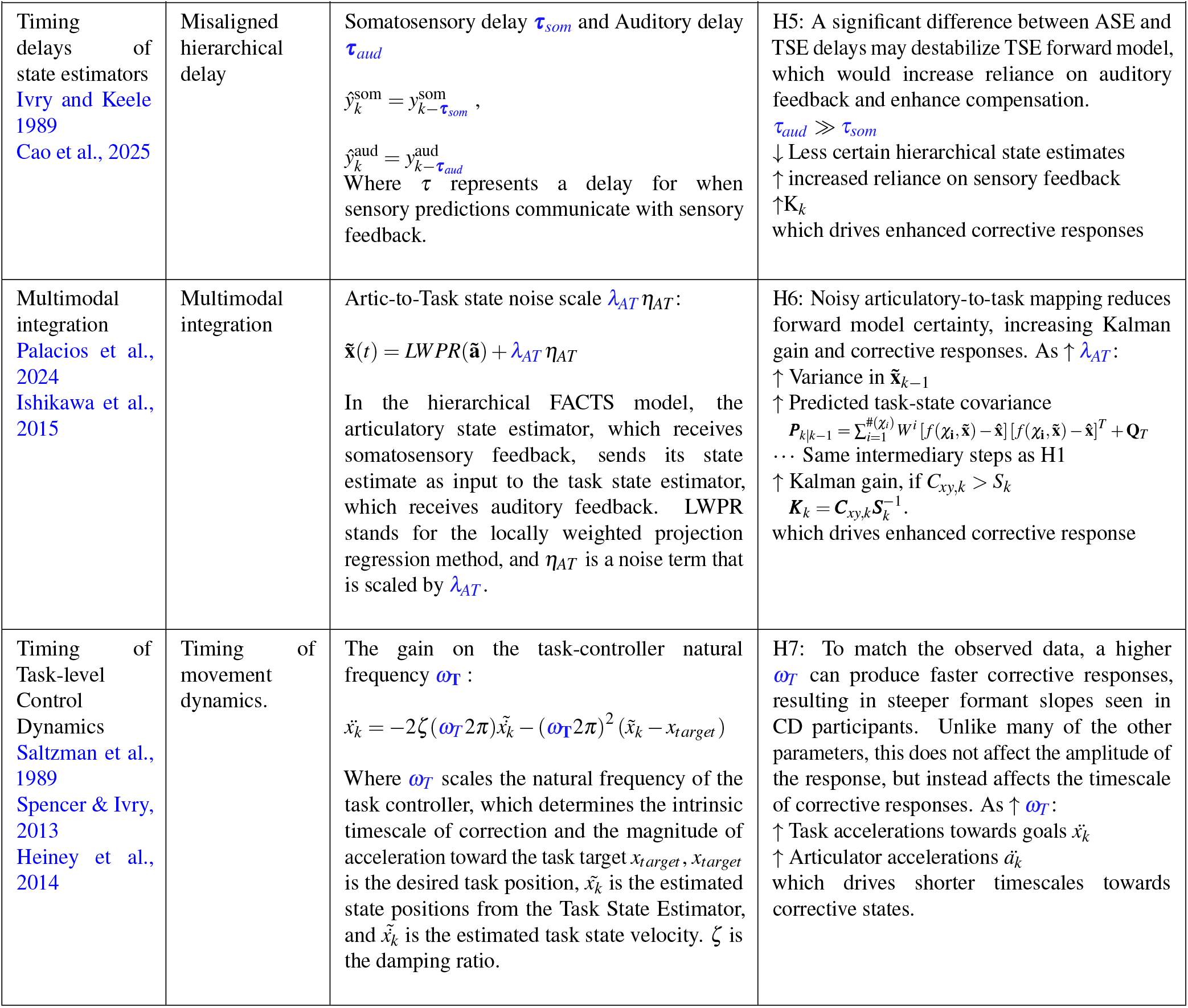
Summary of hypothesized cerebellar mechanisms. Each row corresponds to a theoretical cerebellar framework and the columns correspond to respective computational components, FACTS parameterizations, and a mechanistic analyses for how the parameter may affect speech compensation. The parameters highlighted in blue are the parameters we apply SBI on.

| Cerebellar Framework | Computational Component | FACTS Parameterization | Mechanistic Hypothesis |
| --- | --- | --- | --- |
| Internal modeling<br>Wolpert, Miall, Kawato, 1998<br>Ito, 2008 | Forward Model | Scale Factor for Estimated State Process Noise Scale $\lambda_Q$:<br>$\mathbf{Q}_A = \lambda_{Q_A} \times \begin{bmatrix} Q_{AA} & Q_{A\dot{A}} \\ Q_{\dot{A}A} & Q_{\dot{A}\dot{A}} \end{bmatrix}$ $\mathbf{Q}_T = \lambda_{Q_T} \times \begin{bmatrix} Q_{TT} & Q_{T\dot{T}} \\ Q_{\dot{T}T} & Q_{\dot{T}\dot{T}} \end{bmatrix}$<br>Where $Q_{AA}$ is the covariance of articulator positions, $Q_{\dot{A}\dot{A}}$ is the covariance of the articulator velocities, $Q_{A\dot{A}}$ is the covariance of the articulator positions and velocities, $Q_{TT}$ is the covariance of the task states, $Q_{T\dot{T}}$ is the covariance of the task velocities, and $Q_{\dot{T}\dot{T}}$ is the covariance of the task positions and velocities. | H1: An inflated $\lambda_Q$ increases estimated process noise, which leads to less certain internal forward models, an increased reliance on sensory feedback, and thereby exaggerated compensation. For example, an $\uparrow \lambda_{Q_T}$ leads to:<br>↑ Predicted covariance<br>$\mathbf{P}_{k\vert k-1} = \sum_{i=1}^{\#(Z_i)} W^i [f(\chi_i, \hat{\mathbf{x}}) - \hat{\mathbf{x}}] [f(\chi_i, \hat{\mathbf{x}}) - \hat{\mathbf{x}}]^T + \mathbf{Q}_T$<br>↑ Sigma point spread<br>$\chi_{i,k\vert k-1} = \hat{\mathbf{x}}_{k\vert k-1} \pm \left( \sqrt{(n+\lambda)\mathbf{P}_{k\vert k-1}} \right)$<br>↑ Innovation Covariance<br>$\mathbf{S}_k = \sum_{i=0}^{2n} W_i^{(c)} \left( h(\chi_{i,k\vert k-1}) - \hat{y}_k \right) \left( h(\chi_{i,k\vert k-1}) - \hat{y}_k \right)^T + R_T$<br>↑ Cross covariance<br>$\mathbf{C}_{xy,k} = \sum_{i=0}^{2n} W_i^{(c)} \left[ \chi_{i,k\vert k-1} - \hat{\mathbf{x}}_{k\vert k-1} \right] \left[ Y_{i,k\vert k-1} - \hat{y}_k \right]^T$<br>↑ Kalman gain, if $C_{xy,k} > S_k$<br>$\mathbf{K}_k = \mathbf{C}_{xy,k} \mathbf{S}_k^{-1}$.<br>which drives enhanced corrective responses. |
| | Feedforward Controller (analogous to the Inverse Model) | Articulatory State Feedback Control Law Noise Scale $\lambda_A \eta_A$:<br>$\ddot{a}_k = \mathbf{J}(\tilde{a}_k)^{\dagger} \ddot{x} - \mathbf{J}(\tilde{a}_k)^{\dagger} \mathbf{J}(\tilde{a}, \tilde{a}_k) \ddot{a}_k + \dots$ $(\mathbf{I}_n - \mathbf{J}(\tilde{a}_k)^{\dagger} \mathbf{J}(\tilde{a}_k)) \ddot{a}_d + \ddot{a}_N + \lambda_A \eta_A$<br>Where $\mathbf{J}(\tilde{a}_k)^{\dagger}$ represents the psuedoinverse of the Jacobian, the task-to-articulator mapping. The first term is the desired task-space acceleration mapped to articulator space. The second term corrects for the Jacobian changing over time as the articulators move. The third term accounts for the null-space motion for secondary goals. The fourth term is for intrinsic dynamics. The final term accounts for noise, $\eta_A \sim N(0, 1)$ multiplied by the norm of $\ddot{a}$. $\ddot{a}_k$ is sent to both the plant and the state estimator as an efferent copy. | H2: Although we anticipate $\lambda_A \eta_A$ will have a limited affect on speech compensation because $\ddot{a}$ is sent to both the plant and the state estimator, it is important to describe a potential mechanistic pathway w.r.t the literature. Thereby, an $\uparrow \lambda_A$ leads to:<br>↑ Variance in State Transition<br>$\chi_{k\vert k-1}^{(i)} = f(\chi_{k-1}^{(i)}, \tilde{a}_{k-1})$<br>↑ Predicted covariance<br>$\mathbf{P}_{k\vert k-1} = \sum_{i=1}^{\#(Z_i)} W^i [f(\chi_i, \hat{\mathbf{a}}) - \hat{\mathbf{a}}] [f(\chi_i, \hat{\mathbf{a}}) - \hat{\mathbf{a}}]^T + \mathbf{Q}$<br>↑ Innovation Covariance<br>$\mathbf{S}_k = \sum_{i=0}^{2n} W_i^{(c)} \left( h(\chi_{i,k\vert k-1}) - \hat{y}_k \right) \left( h(\chi_{i,k\vert k-1}) - \hat{y}_k \right)^T + R_T$<br>↑ Cross covariance<br>$\mathbf{C}_{xy,k} = \sum_{i=0}^{2n} W_i^{(c)} \left[ \chi_{i,k\vert k-1} - \hat{\mathbf{a}}_{k\vert k-1} \right] \left[ Y_{i,k\vert k-1} - \hat{y}_{i,k\vert k-1} \right]^T$<br>↑ Kalman gain, if $C_{xy,k} > S_k$<br>$\mathbf{K}_k = \mathbf{C}_{xy,k} \mathbf{S}_k^{-1}$.<br>which drives enhanced corrective responses. |
| Error Detection & Correction<br>Ito, Marrs, Albus 1970s | Observation uncertainty in auditory prediction. | Auditory Noise Scale $\lambda_{R_T} R_T$. The Kalman gain can be expanded:<br>$\mathbf{K}_k = \tilde{\mathbf{X}} \mathbf{W} \tilde{\mathbf{Y}}^T \left( \tilde{\mathbf{Y}} \mathbf{W} \tilde{\mathbf{Y}}^T + \lambda_{R_T} \mathbf{R}_T \right)^{-1}$<br>Where $\mathbf{R}_T$ is the observation covariance, $\lambda_{R_T}$ scales the observation covariance, $\tilde{\mathbf{Y}} \mathbf{W} \tilde{\mathbf{Y}}^T$ is the covariance of observations, and $\tilde{\mathbf{X}} \mathbf{W} \tilde{\mathbf{Y}}^T$ is the cross covariance between state and observation estimates. Uncertainty in observations affects certainty in errors, and can intensify corrective responses. | H3: As $\downarrow \lambda_{R_T}$:<br>↓ Innovation Covariance<br>$\mathbf{S}_k = \tilde{\mathbf{Y}} \mathbf{W} \tilde{\mathbf{Y}}^T + \lambda_{R_T} \mathbf{R}_T$<br>$= \sum_{i=0}^{2n} W_i^{(c)} \left( h(\chi_{i,k\vert k-1}) - \hat{y}_k \right) \left( h(\chi_{i,k\vert k-1}) - \hat{y}_k \right)^T + \lambda_{R_T} R_T$<br>↑ Kalman Gain<br>$\mathbf{K}_k = \mathbf{C}_{xy,k} \mathbf{S}_k^{-1}$<br>which drives enhanced corrective responses. |
| | Observation uncertainty in somatosensory prediction. | Somatosensory Noise Scale $\lambda_{R_A} R_A$. Similar to Auditory noise scale, except in the Articulatory State Estimator.<br>$\mathbf{K}_k = \tilde{\mathbf{X}} \mathbf{W} \tilde{\mathbf{Y}}^T \left( \tilde{\mathbf{Y}} \mathbf{W} \tilde{\mathbf{Y}}^T + \lambda_{R_A} \mathbf{R}_A \right)^{-1}$<br>Where $\mathbf{K}_k = \mathbf{C}_{xy,k} \mathbf{S}_k^{-1}$ is expanded into its matrix compoments. | H4: As $\downarrow \lambda_{R_A}$:<br>Similar to $\downarrow \lambda_{R_T}$, except discrepancies are accounted for in the somatosensory state space. Since task perturbations are in auditory state space, we do not anticipate this parameter will have much effect. |

**Table 1.** Summary of hypothesized cerebellar mechanisms.
| Cerebellar Theoretical Framework | Computational Component | FACTS Parameterization | Mechanistic Hypothesis |
| --- | --- | --- | --- |
| Timing delays of state estimators<br>Ivry and Keele 1989<br>Cao et al., 2025 | Misaligned hierarchical delay | Somatosensory delay $\tau_{som}$ and Auditory delay $\tau_{aud}$<br>$\hat{y}_k^{som} = y_{k-\tau_{som}}^{som}$<br>$\hat{y}_k^{aud} = y_{k-\tau_{aud}}^{aud}$<br>Where $\tau$ represents a delay for when sensory predictions communicate with sensory feedback. | H5: A significant difference between ASE and TSE delays may destabilize TSE forward model, which would increase reliance on auditory feedback and enhance compensation.<br>$\tau_{aud} \gg \tau_{som}$<br>↓ Less certain hierarchical state estimates<br>↑ increased reliance on sensory feedback<br>↑ $K_k$<br>which drives enhanced corrective responses |
| Multimodal integration<br>Palacios et al., 2024<br>Ishikawa et al., 2015 | Multimodal integration | Artic-to-Task state noise scale $\lambda_{AT} \eta_{AT}$:<br>$\tilde{\mathbf{x}}(t) = LWPR(\tilde{\mathbf{a}}) + \lambda_{AT} \eta_{AT}$<br>In the hierarchical FACTS model, the articulatory state estimator, which receives somatosensory feedback, sends its state estimate as input to the task state estimator, which receives auditory feedback. LWPR stands for the locally weighted projection regression method, and $\eta_{AT}$ is a noise term that is scaled by $\lambda_{AT}$. | H6: Noisy articulatory-to-task mapping reduces forward model certainty, increasing Kalman gain and corrective responses. As $\uparrow \lambda_{AT}$:<br>↑ Variance in $\tilde{\mathbf{x}}_{k-1}$<br>↑ Predicted task-state covariance<br>$\mathbf{P}_{k\vert k-1} = \sum_{i=1}^{\#(\mathcal{Z}_i)} W^i [f(\chi_i, \tilde{\mathbf{x}}) - \hat{\mathbf{x}}] [f(\chi_i, \tilde{\mathbf{x}}) - \hat{\mathbf{x}}]^T + \mathbf{Q}_T$<br>... Same intermediary steps as H1<br>↑ Kalman gain, if $C_{xy,k} > S_k$<br>$\mathbf{K}_k = \mathbf{C}_{xy,k} \mathbf{S}_k^{-1}$.<br>which drives enhanced corrective response |
| Timing of Task-level Control Dynamics<br>Saltzman et al., 1989<br>Spencer & Ivry, 2013<br>Heiney et al., 2014 | Timing of movement dynamics. | The gain on the task-controller natural frequency $\omega_T$:<br>$\ddot{x}_k = -2\zeta(\omega_T 2\pi)\dot{x}_k - (\omega_T 2\pi)^2(\tilde{x}_k - x_{target})$<br>Where $\omega_T$ scales the natural frequency of the task controller, which determines the intrinsic timescale of correction and the magnitude of acceleration toward the task target $x_{target}$, $x_{target}$ is the desired task position, $\tilde{x}_k$ is the estimated state positions from the Task State Estimator, and $\dot{\tilde{x}}_k$ is the estimated task state velocity. $\zeta$ is the damping ratio. | H7: To match the observed data, a higher $\omega_T$ can produce faster corrective responses, resulting in steeper formant slopes seen in CD participants. Unlike many of the other parameters, this does not affect the amplitude of the response, but instead affects the timescale of corrective responses. As $\uparrow \omega_T$:<br>↑ Task accelerations towards goals $\ddot{x}_k$<br>↑ Articulator accelerations $\ddot{a}_k$<br>which drives shorter timescales towards corrective states. |

Inferences about underlying neural mechanisms rely on the ability to reproduce behavior associated with both normal and disordered brains using computational models (e.g., [44, 33, 34, 35, 45]); however, both parameterizing neural computations that drive participant behavior and estimating certainty over parameter values has historically been challenging. Modern approaches such as Simulation Based Inference (SBI) offers a principled approach [46, 47] to resolve this ambiguity in mechanistic function. SBI integrates computational modeling and Bayesian inference and, when paired with hypothesis testing, can reveal which parameters in the computational model best explain behavioral differences between two different sets of data. For instance, previous work, such as Gaines et al., [48], used SBI to disambiguate CD vs. HC mechanisms of enhanced vocal pitch compensation. Our work extends that study by using a more complex model of speech motor control, expanding the set of hypotheses tested, and investigating a difference speech task.

The prior work of Gaines et al. [48] demonstrated the effectiveness of SBI to estimate control parameters underlying pitch compensation behavior, setting the groundwork that this work extends. In Gaines et al., pitch was controlled using a state feedback control (SFC) model ([45]) that controlled pitch via a simple spring-mass system in which the length of the spring represents muscle tension on the vocal folds. With their SFC model, the authors found that pitch compensation may be driven by both the feedback noise ratio and the controller gain. However, the SFC model does not include task dynamics ([49]), and thus could not model the shape of the vocal tract and thereby the resonances of voice such as F1, F2, F3, etc. Most importantly, their simplified control architecture limited the range of hypotheses about cerebellar (dis)function that could be tested with SBI. To address the broader range of hypotheses, here we use the Hierarchical Feedback-Aware Control of Tasks in Speech (FACTS) model [33, 34].

FACTS is a relatively newer model of speech motor control that combines the strengths of both state feedback control [45] and task dynamics [49]. FACTS contains nested controllers relating to high-level tasks (control of vocal tract constrictions) and low-level articulation (control of jaw, lips, and tongue). With the ability to investigate aspects of speech beyond pitch, this work investigated compensatory responses to the first vowel format (F1). F1 is the lowest resonance of the vocal tract, shifting it changes the way vowels sound [50, 51]. Critically, this more complex control model allows us to investigate a broader range of potential cerebellar functions underlying the observed increase in online compensatory F1 response in adults with CD.

Here, we pair FACTS with SBI to test if any of the five proposed cerebellar functions (parameterized nine ways) in motor control can explain the observed differences in F1 compensation. We also attempt to disentangle the relative contributions between the HC and CD groups. The mechanisms we test are summarized in Table 1: internal modeling [5, 12], error detection and correction [15, 16, 14], delay processing [22, 24], multi-modal integration [18], and timing of movement dynamics [52]. For each proposed mechanism, we identify the parameters of the FACTS model associated with the proposed computational process (Table 1; Columns 2-3). We then illustrate how impairments in each of these proposed mechanisms could lead to the observed increase in F1 compensation in individuals with CD (Table 1; Column 4). Together, Table 1 provides the theoretical-to-model mapping used for inference.

We briefly summarize that mapping here to clarify how each proposed cerebellar computation is operationalized within FACTS. The certainty of the forward model, a type of internal model that predicts the outcomes of actions, is related to the parameters 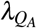 and 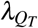, which scale the noise in the covariance matrices used to estimate the (lower level) articulatory and (higher level) task states, respectively. The articulatory state estimator uses the covariance of articulator positions and velocities; the task state estimator uses the task (i.e. constriction degree) position and velocities. While FACTS does not include an inverse model per se, a similar function is instantiated with the articulatory feedforward controller; here, the parameter *λ*_*A*_ scales the noise in the articulatory state feedback control signal. Error detection and correction are reflected by the parameters 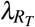 and 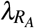, which affect the observation uncertainty in auditory and somatosensory systems, respectively, by scaling each of the associated observation covariance matrices. Delay processing, reflected by sensory processing delays denoted by the parameters *τ*_*som*_ and *τ*_*aud*_, models the length of delay in the articulatory and task state estimators, respectively. Multi-modal integration is reflected by *λ*_*AT*_, which scales the amount of noise in the transformation between articulatory and task state estimates. Finally, the timing of movement dynamics is operationalized as a gain on the natural frequency of the task controller, denoted by *ω*_*T*_ . This parameter modulates the damping and velocity terms of the damped spring mass equation, and thereby affects the timescale of corrections towards a goal state.

By testing multiple proposed cerebellar functions within a single modeling and inference pipeline, we move beyond isolated assessment of mechanisms underlying behavioral changes in CD to instead simultaneous assessment that can uncover interacting, joint structure of model mechanisms. This approach, illustrated in Figure 1, provides an interpretable link between theories of cerebellar (dis)function and behavioral observations. This approach also supports a potential mechanistic characterization of the changes in speech behavior observed in CD, and provides a generalizable framework for mechanistic disambiguation.

**Figure 1.**
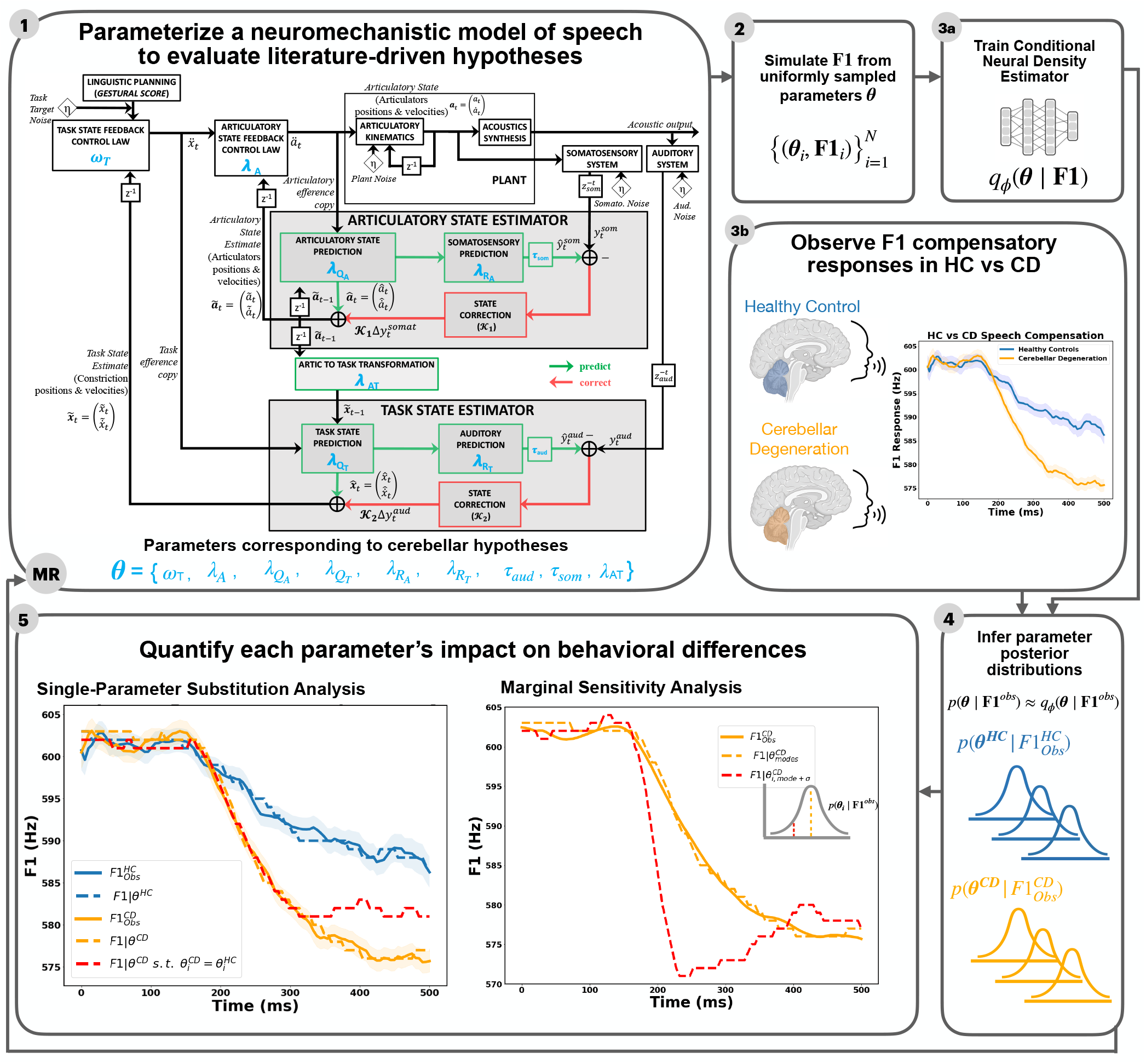
Summary of research paradigm. (1) The FACTS model implements hierarchical state feedback control of speech based on articulatory- and task-level states. This model includes tunable parameters *θ* that correspond to hypothesized cerebellar functions to sensorimotor prediction and control. (2) We simulate speech formants **F1** sampling from a uniform prior, generating a dataset 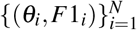, for *N* = number of simulations. (3a) A conditional neural density estimator *q*_*φ*_ (*θ*|*F*1) is trained to estimate parameters *θ* given *F*1. (3b) Empirical speech differences in HC vs. CD groups are observed. (4) Given empirical observations of group averaged healthy control 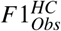 or cerebellar degeneration 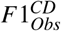, we infer respective posterior distributions over parameters 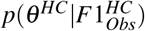 and 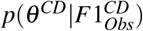. (MR) Multi-Round SBI can be performed for fine-tuning by using the posterior of the current round as the proposal for the next round. (5) Finally, we analyze how individual parameters contribute to differences in behavior using single-parameter substitution, marginal sensitivity, and posterior distribution analyses.

## Results

We combined a hierarchical mechanistic model of speech motor control (FACTS) with simulation-based inference to infer control mechanism differences in online F1 compensatory responses to auditory F1 perturbations between CD 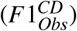 vs HC 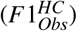 groups. Using the FACTS model, we simulated 1,002,080 speech trajectories by uniformly sampling a nine-dimensional parameter space corresponding to hypothesized cerebellar computations. We then trained a conditional neural density estimator on this set of parameter vectors and F1 trajectories {*θ, F*1} ^*N*=1,002,080^. To fine-tune the conditional density estimator, several rounds of multi-round inference were trained using 500 training samples with the posterior in the previous round serving as the proposal in the current round. We then inferred parameter posterior distributions conditioned on empirical 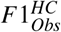 and 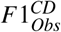, and evaluated model fidelity using posterior predictive checks. We quantified the contribution of these parameter estimates to evaluate the differences between CD and HC behavior in four ways: Glass Δ effect sizes between parameter posteriors for the two groups; single-parameter substitution analysis swapping each parameter, one at a time, in the estimated CD set with the equivalent parameter in the estimated HC set; marginal sensitivity analysis for each parameter estimate; and joint parameter posterior analysis. Together these analyses allowed us to assess which computational components best explain observed CD vs. HC differences in F1 compensation.

### SBI with FACTS accurately recapitulates speech behavior and group differences

To test that inference under the FACTS–SBI framework produces reliable results, we performed posterior predictive checks (PPC) [53, 54]. The respective modes of the marginal posterior distributions of the FACTS model parameters for HC and CD were used to generate ten F1 trajectories for each group, the average of which was compared with the average empirical F1 data. Mean FACTS model trajectories generated with SBI-derived parameter estimates closely matched the temporal dynamics of the empirical compensatory behavior for both HC and CD groups (Figure 2).

**Figure 2.**
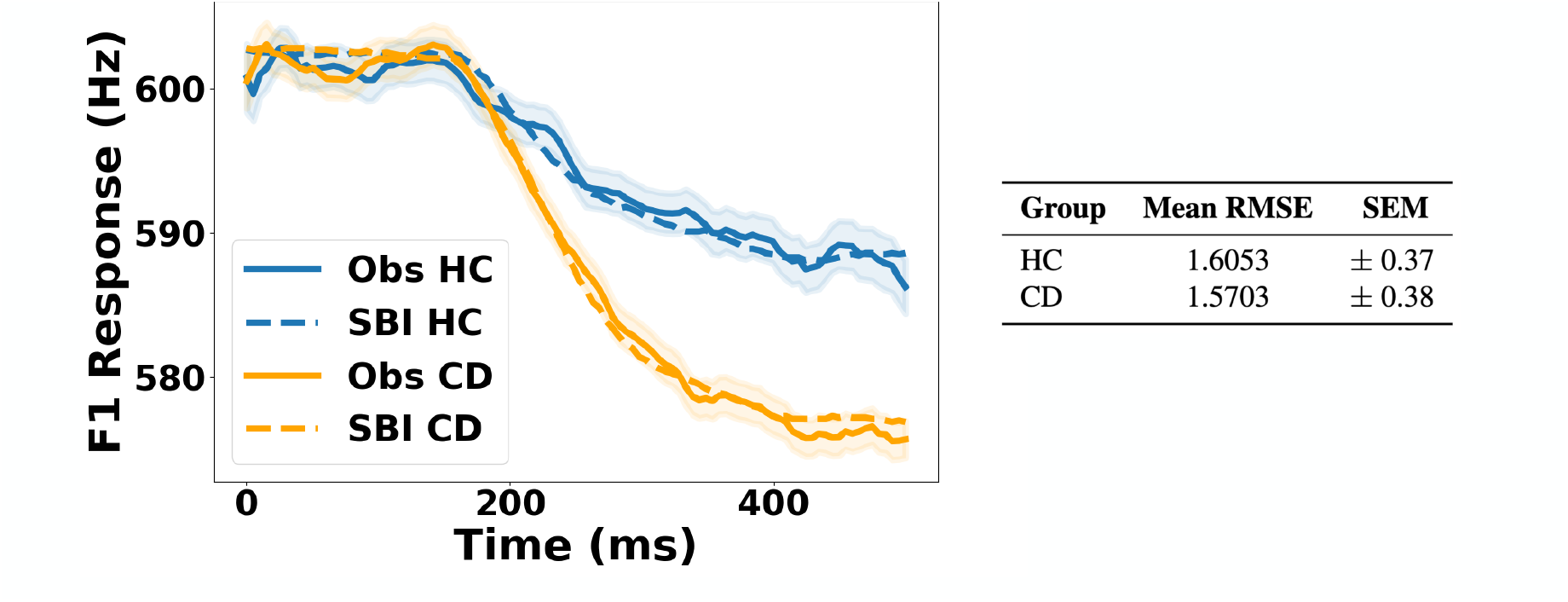
Left: Observed group mean F1 trajectories (solid) from Parrell et al. (2017). SBI Simulated trajectories in FACTS using SBI-estimated parameters for each group (dashed), yield close agreement in both response magnitude and temporal dynamics for both HC (blue) and CD (orange) groups. Shaded bands denote variability around the observed behavioral F1 trajectories (SEM). The table on the right reports RMSE and SEM between average model and observed F1 trajectories.

### The CD group differs in internal modeling, timing of movement dynamics, and multimodal integration mechanisms - marginal posterior distribution analysis

To identify which inferred mechanisms most strongly separated healthy controls (HC) from individuals with cerebellar degeneration (CD), we calculated Glass’s Δ effect sizes between HC and CD groups for each parameter’s marginal posterior distribution (Figure 3). The process-noise scale of the task-state forward model 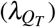, reflecting the (un)certainty of the forward model for task-level control, had the largest Glass’s |Δ| with +30.69. The timing of movement dynamics *ω*_*T*_ had the second largest Glass’s |Δ| of -1.87. The noise added between the multi-modal, articulatory-to-task transformations (*λ*_*AT*_ ) had a Glass’s |Δ| of 0.81. The remaining parameters had Glass’s |Δ| below 0.50.

**Figure 3.**
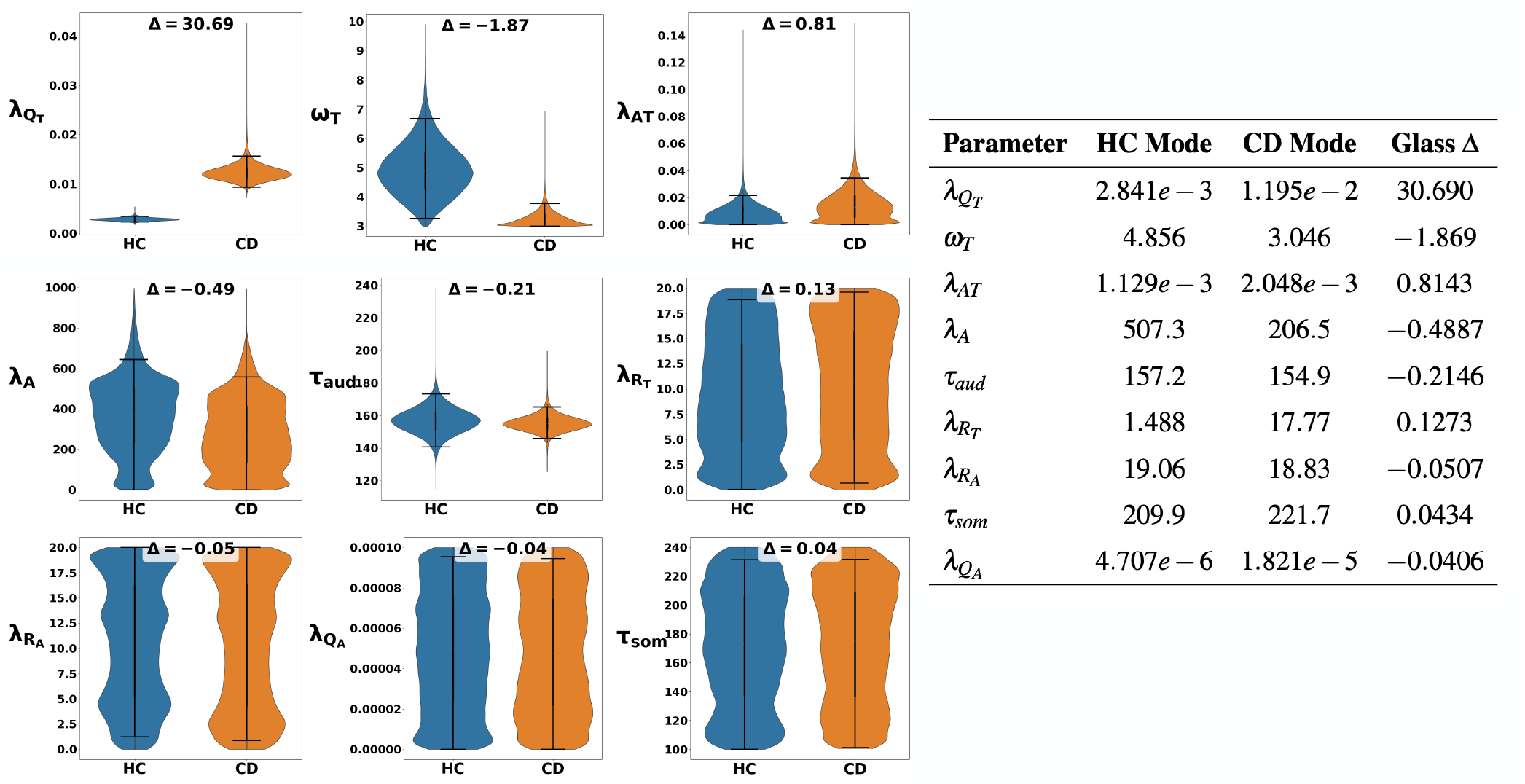
Left: violin plots of the marginal posterior distributions for each parameter comparing the HC (blue) and CD (orange) groups, based on 100,000 posterior samples drawn from the trained neural posterior density estimator conditioned on the observed data. Highest Density Intervals (black lower- and upper-bound horizontal lines) show the narrowest credible interval containing a 95% probability mass. Right: Posterior modes, and Glass Effect Sizes for different parameters. The three parameters with effect sizes > 0.5 are 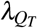, *ω*_*T*_, and *λ*_*AT*_ .

### Internal forward modeling and timing of movement dynamics mechanisms drive differences in CD formant compensation control - single parameter substitution analysis

To test which parameters play the most important role in driving the difference between compensation in the HC vs. CD groups, we performed single-parameter substitution analysis for each parameter. For this analysis, we substituted each estimated parameter for the CD group, one at a time, with the estimated value for the HC group, while keeping the remaining parameters fixed to their estimated posterior modes (Figure 4). We quantified the effect of each substitution by comparing its model fit (RMSE) of FACTS F1 trajectories to the fit of the baseline FACTS behavior (i.e. where all parameters are fixed to their posterior modes). Twenty-five F1 simulations were generated with each parameter set so means and variances could be calculated. The results of this analysis showed that the most important parameter driving the between-group difference was the task forward model’s process noise scale 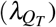, with a large increase in the model-fit RMSE relative to the baseline (CD Baseline= 1.727*±*0.084; 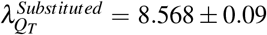; Glass Δ = 31.031; *p*_*MWU*_ = 1.42 *×* 10^−9^). A smaller but still substantial effect was found for the timing of movement dynamics timescale parameter (*ω*_*T*_ ), (CD Baseline= 1.803*±*0.132; 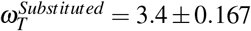; Glass Δ = 9.892; *p*_*MWU*_ = 1.42 *×* 10^−9^). The remaining parameters have smaller impact with less than 1 RMSE difference compared to baseline, and absolute values of Glass delta < 1.0.

**Figure 4.**
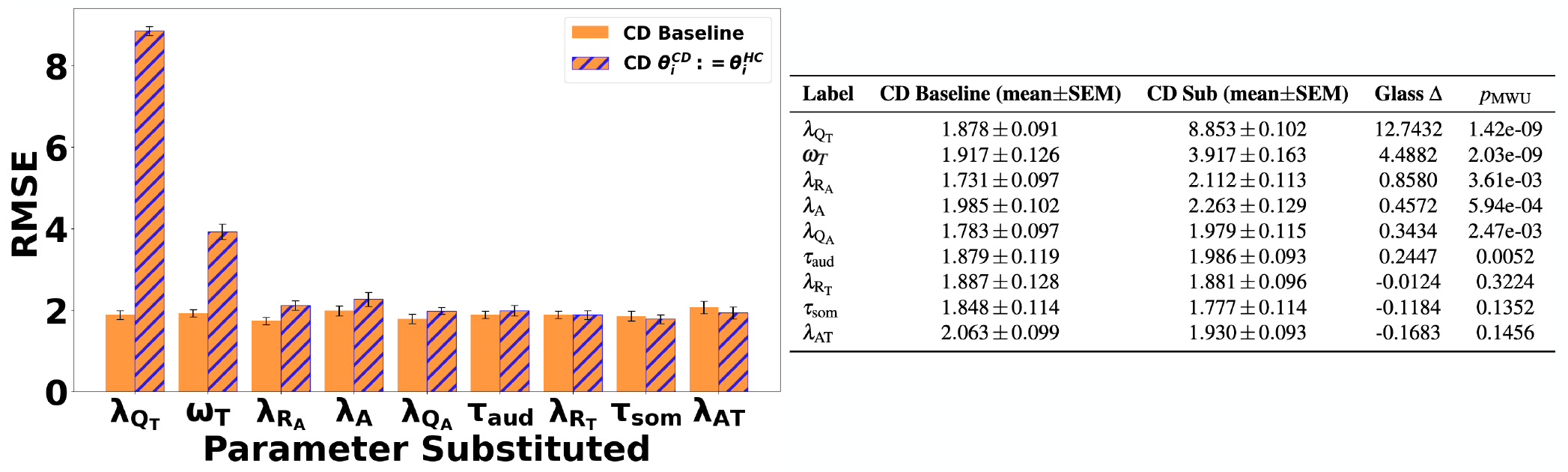
Left: Results of the substitution analysis where each estimated model parameter for the CD group was swapped, one-at-at-time, with the equivalent estimated parameter from the HC group (orange with blue hatches), compared to the baseline model with all CD-specific parameter estimates (orange). RMSE reflects the error between the FACTS trajectories generated with each model and the behavioral CD data. Each condition was simulated 25 times to estimate mean ± SEM (error bars). Right: Table showing each parameter’s Baseline RMSE (mean ± SEM), Substituted RMSE (mean ± SEM), Glass’s Δ, and Mann–Whitney U p-values based on 25 substitution rounds per parameter.

### Internal forward modeling and timing of movement dynamics impact model output variance the most - marginal sensitivity analysis

Although previous analyses identified key drivers of CD–HC behavioral differences, it remained unclear how small deviations around the posterior mode of each parameter influence simulated F1 trajectories. To determine how changes in each control parameter affects the F1 speech trajectory, we conducted a marginal sensitivity analysis on the estimated FACTS parameters. Figure 5 illustrates how much FACTS output changed, for both HC and CD groups, when a single input parameter was varied around its estimated mode, while holding all other parameters constant. Parameters that led to the most behavioral difference for both groups were the task forward model process noise scale 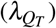 and the natural frequency of the task controller (*ω*_*T*_ ), with baseline RMSE approximately at 2 Hz *±*0.1 Hz, whereas large deviations of *±*3*σ* led to RMSE ≥ 2.5 Hz *±*0.1. The task forward model process noise scale 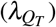 affected both the speed and amplitude of corrective responses, while the natural frequency of the task controller (*ω*_*T*_ ) inversely affected the timescale of corrective responses. Other parameters showed moderate sensitivity: large deviations from the mode *±*3*σ* showed an increase in RMSE error, but below the 2.5Hz threshold. Those parameters include the delay applied to the task-state-estimator auditory predictions (*τ*_*aud*_), which does not change the amplitude of compensation, but changed the response latency; the multi-modal integration noise (*λ*_*AT*_ ); and the noise added to the feedforward controller (*λ*_*A*_). Varying the remaining parameters showed relativity minimal effects on F1 output compared to baseline.

**Figure 5.**
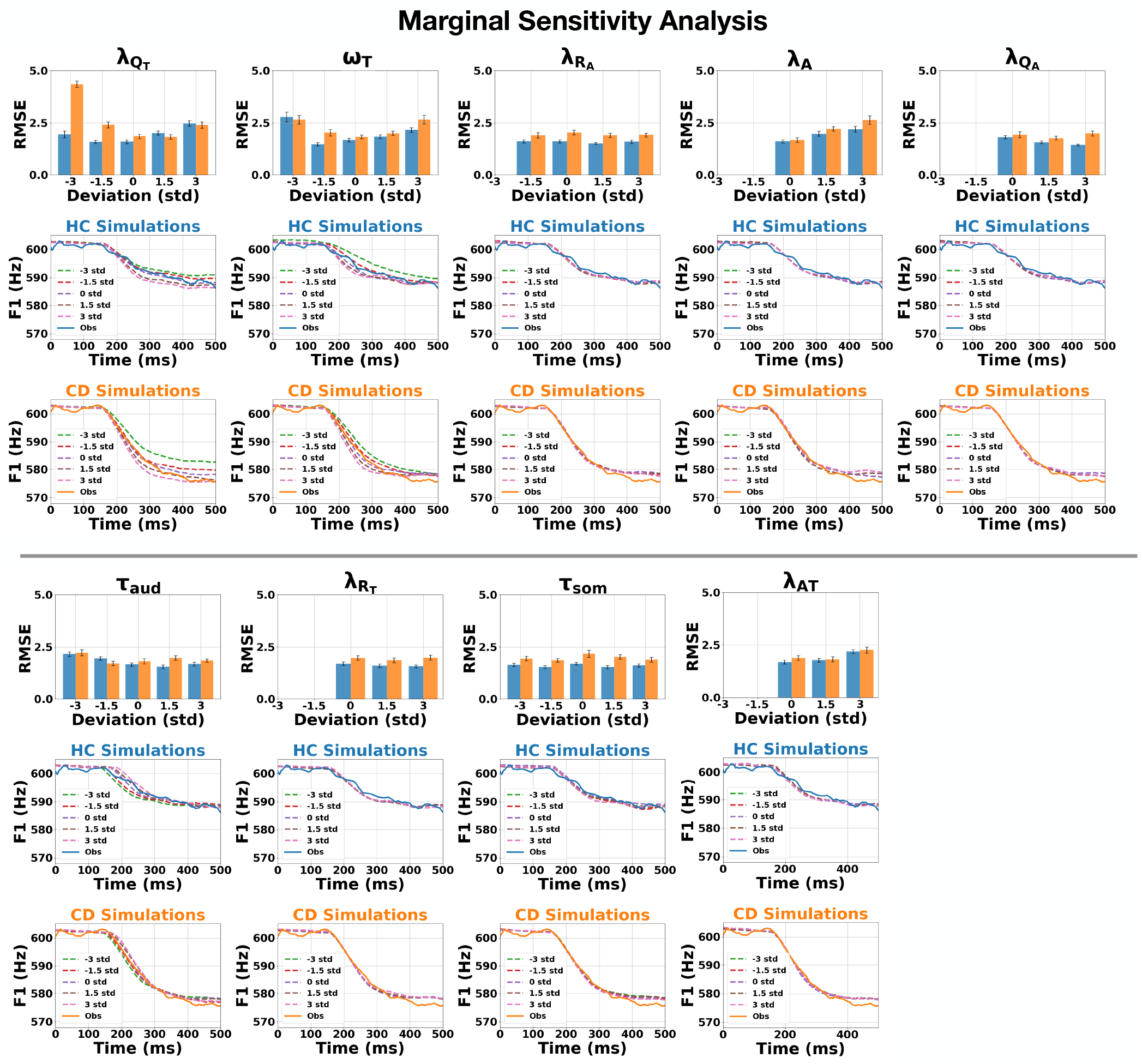
Results of the marginal sensitivity analysis for each estimated parameter for HC (blue) and CD (orange) groups. The first row displays changes in model fit (RMSE) when varying the parameter of interest around its estimated mode while freezing all parameter values to their estimated modes. The second row displays compensatory F1 responses for the HC group, showing empirical data in solid lines and simulation data in dotted lines (color reflects varying parameter values) The third row displays data for the CD group, in the same manner as the row above. For subplots where some std values show no barplots, the parameter value returned negative values and no simulations were run.

### Joint posterior distributions reveal tradeoffs between (Timing of Movement Dynamics vs. Forward Model) and (Timing of Movement Dynamics vs. Multi-Modal Integration) mechanisms

There were two findings from our results that motivated a joint parameter posterior analysis. The first result is the slower timing of movement dynamics (*ω*_*T*_ ) in CD than what was hypothesized (H7; Table 1). The second result was the multi-modal integration noise (*λ*_*AT*_ ) having a large posterior Glass Δ, but showing less of a contribution in the single-parameter substitution analysis. These results either contradicted what we expected, or the analysis introduced more questions. In general, the results highlighted a limitation with the previous analyses: the analyses change parameters one at a time, while keeping other parameters constant (i.e. ceteris paribus). Although ceteris paribus analyses can isolate variables and make results easier to understand, they fail to reveal parameter regimes of joint structure, synergies, and trade-offs that give a more holistic view of multi-mechanism interactions. To extend beyond this limitation, we investigated parameter joint interactions to provide better explanations.

The results prior showed that the main differences in model parameters that drive HC vs. CD compensation differences are, in order of relative importance, the task forward model process noise scale 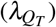, timing of movement dynamics (*ω*_*T*_ ), and multi-modal integration (*λ*_*AT*_, though this latter parameter was not as consistently identified across the three analyses). Here, we focus on how these parameters may covary by plotting the joint posterior distributions between each pair of these three parameters, and quantifying their covariation. We quantify their covariation in three ways: Spearman’s *ρ*, mutual information (MI), and Hellinger distance, which measure complementary aspects of posterior dependence. Spearman’s *ρ* captures the direction and monotonic strength of pairwise dependence. Mutual information measures how the variables jointly resolve ambiguity even for non-linear relationships. Hellinger distance measures the overall discrepancy between the joint posterior vs the product of marginals which captures degree of dependence. Larger absolute values of *ρ*, larger mutual information values, and larger Hellinger distances all indicate stronger pairwise posterior dependence and thereby suggest mechanistic dependencies and potential trade-offs or synergies. These values in conjunction with the geometry of the joint posterior plots illustrate the degree, direction, and boundaries of control mechanism interaction, and thereby quantifies their level of (dis)entanglement.

Joint posterior dependence analysis revealed dependence for two out of the three pairs of parameters. For the (timing of movement dynamics vs. forward model) pair (*ω*_*T*_, 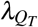), posterior dependence was negative in both CD and HC, with stronger dependence in healthy controls (HC; Spearman’s *ρ* = −0.494, mutual information = −0.160 nats, Hellinger distance = 0.207) than in cerebellar degeneration (CD; *ρ* = −0.325, mutual information = 0.077 nats, Hellinger distance = 0.139). The negative, diagonal ridges in the geometry of the joint posterior plots illustrate probability contours where increasing the value of one parameter, and decreasing the value of the other parameter, may lead to the same behavioral output (i.e. a negative trade-off regime).

A similar pattern was observed for the (timing of movement dynamics vs. multi-modal integration) pair (*ω*_*T*_, *λ*_*AT*_ ). Posterior dependence was again negative in both groups and stronger in HC (*ρ* = −0.463, mutual information = 0.150 nats, Hellinger distance = 0.199) than in CD (*ρ* = −0.272, mutual information = 0.063 nats, Hellinger distance = 0.124). This also suggests a negative trade-off with the timing of movement dynamics *ω*_*T*_ again being a key component.

In contrast, the (forward model vs. multi-modal integration) pair (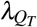, *λ*_*AT*_ ) exhibited comparatively weak dependence in both HC and CD. In CD, posterior Spearman correlation was near zero (*ρ* = 0.004), with low mutual information (0.013 nats) and low Hellinger distance (0.056). In HC, dependence was slightly larger but remained weak (*ρ* = 0.127, mutual information = 0.025 nats, Hellinger distance = 0.075).

Taken together, these results show that pairwise posterior structure was present for both (timing of movement dynamics vs. forward model) and (timing of movement dynamics vs. multi-modal integration) with negative sloped trade-offs. In contrast, dependence between multi-modal integration (*λ*_*AT*_ ) and forward model 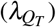 mechanisms was less strong. Across all pairs, all metrics indicated stronger posterior coupling in HC than in CD. The interpretation of these results are elaborated on in the discussion section.

## Discussion

Although behavioral studies have shown that individuals with cerebellar degeneration often produce exaggerated compensatory responses to auditory perturbations, the underlying mechanisms that drive this difference in behavior remains unresolved. One challenge in identifying the mechanisms is the large number of complementary theories of cerebellar function in motor control. These theories include the cerebellum’s role in internal modeling [9, 7, 8, 12, 13], sensory-error processing [14, 15, 16], multimodal sensory integration [17, 18, 19, 20, 21], delay processing [22, 23, 24], and the timing of movement dynamics [25, 26, 27, 28]. Our study helps resolve this gap between behavior and underlying control impairments. This work demonstrates that simulation-based inference can infer probabilities over the control parameters of a computational model of speech production (FACTS) to not only reproduce group-level behavioral differences between HC and CD, but crucially also quantify relative contributions of control mechanisms. While it is important to note these analyses do not necessarily disprove any specific hypothesis, they nonetheless serve to suggest the relative importance of different control mechanisms under specific modeling, parameterization, and task assumptions.

### Mechanistic Contribution of the Cerebellum to Error Correction

Our results largely support the view that cerebellar degeneration impairs internal predictive models [55], and thereby increases the gain on sensory feedback control. Among all parameters examined, the forward model process noise scale 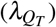 of the task-state estimator consistently emerged as the strongest differentiator between the HC and CD groups, including in direct estimates of effect size, the single-parameter substitution analysis, and in the marginal posterior analysis. Our results suggest that the increase in the forward model process noise scale 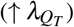 of the CD group leads to less certain forward models, which in turn drives an increased gain on sensory feedback in the Kalman filter underlying task-state estimation, and ultimately an exaggerated corrective response to the auditory feedback perturbation. These results support theoretical accounts that position the cerebellum as a locus of forward-modeling [56], where increased uncertainty in prediction may lead to stronger sensory-driven corrections [48].

Our results also support the view that cerebellar degeneration impairs timing mechanisms associated with movement dynamics [23, 27, 52]. Contrary to this investigation’s initial hypothesis (H7; Table 1) that the timing of movement dynamics parameter would be faster (↑*ω*_*T*_ ) in the CD group compared to HC because CD showed faster compensation behavior, our SBI results instead showed the CD group had a slower timing of movement dynamics value (↓*ω*_*T*_ ), which is consistent with at least one prior report [57]. Hypothesis H7 predicted CD to have faster timing of movement dynamics than HC 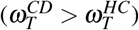 because the F1 compensation velocity of CD is greater than HC 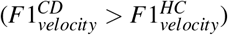. Indeed, the Marginal Sensitivity Analysis (Figure 5, column *ω*_*T*_ ), shows results consistent with H7’s mechanistic analysis where greater *ω*_*T*_ resulted in faster F1 trajectories. However, our SBI-derived results showed the opposite, 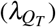; Glass’s Δ = −1.896). In other words, even though the CD speech behavior is faster, the CD group nonetheless shows slower timescale dynamics in their task-controller to correct towards goal states. In an attempt to explain this result, we anticipated there may be an interaction among several parameters that trade-off to produce the observed behavior. We hypothesized that if the forward model process noise scale 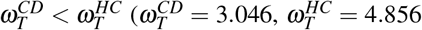 drives most of the compensatory F1 behavior in CD, then the gain on timing of movement dynamics (*ω*_*T*_ ) may serve to counteract and potentially balance F1 compensations. If this were true, we are likely to see a trade-off geometry in the joint posterior distributions. To more closely investigate this, we performed joint parameter posterior analysis, and discuss the results in the later half of the discussion section. Nonetheless, our results support the hypothesis [52, 26] that cerebellar timing computations are a major contributor of motor behavior and online corrective responses.

Our results also support the view that cerebellar degeneration impairs multi-modal integration [17, 19, 20]. The multi-modal integration parameter (*λ*_*AT*_ ) showed the third largest impact in the marginal parameter posterior analysis (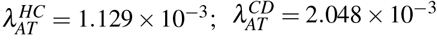; Glass Δ = 0.8143), but did not show a significant difference in the single-parameter substitution analysis. *λ*_*AT*_ is the noise added to the transformation between the articulatory state estimator and the task state estimator. Our findings suggest that *λ*_*AT*_ may differ systematically between CD and HC at the level of the inferred marginal posterior, but is unlikely to act as a primary independent driver of the behavioral differences in the model. That is, the CD group may have more noise in its multi-modal integration calculations, but multi-modal integration is not the primary driver of the enhanced compensatory F1 behavior. Thus, our results partially support cerebellar accounts of multimodal integration and cross-domain state coupling [17, 18]. A joint posterior analysis may give insight as to why there is a marginal posterior difference for this mechanism, but not a difference in this mechanism driving behavior.

Two discrepancies motivated the joint posterior analysis. The first discrepancy is the slower timing of movement dynamics (↑*ω*_*T*_ ) in CD than what was hypothesized (↓*ω*_*T*_ ; H7; Table 1). The second discrepancy was the multimodal integration noise (*λ*_*AT*_ ) resulting in a large posterior Glass Δ, but showing less of a contribution to speech behavior in the single-parameter substitution analysis. If the parameters in isolation reveal counter-intuitive results, then their joint interactions may provide better explanations.

The negative posterior dependence between the timing of movement dynamics and forward-model parameters provides one such explanation (Figure 6; 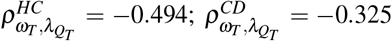). Across posterior solutions, larger values of the forward-model parameter 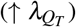 tended to co-occur with smaller values of the movement-dynamics parameter (↓*ω*_*T*_ ). CD exhibited this joint pattern: substantially larger 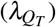 values (Glass Δ = 30.69) accompanied by smaller (*ω*_*T*_ ) values (Glass Δ = ™1.87). Thus, the decrease in (*ω*_*T*_ ) should not necessarily be interpreted as a strict contradiction of H7. Instead, it appears to form part of a joint parameter regime in which greater noise in the forward model is associated with slower movement dynamics. This interpretation is consistent with reports of slower corrective dynamics in cerebellar degeneration [57] and impaired forward modeling following cerebellar damage [58]. Although our results corroborate these accounts in the literature, they discussed their results as isolated mechanisms of cerebellar dysfunction. In contrast, our work extends prior studies by providing a quantitative characterization – including the direction, mutual information, and strength – of mechanism interactions.

**Figure 6.**
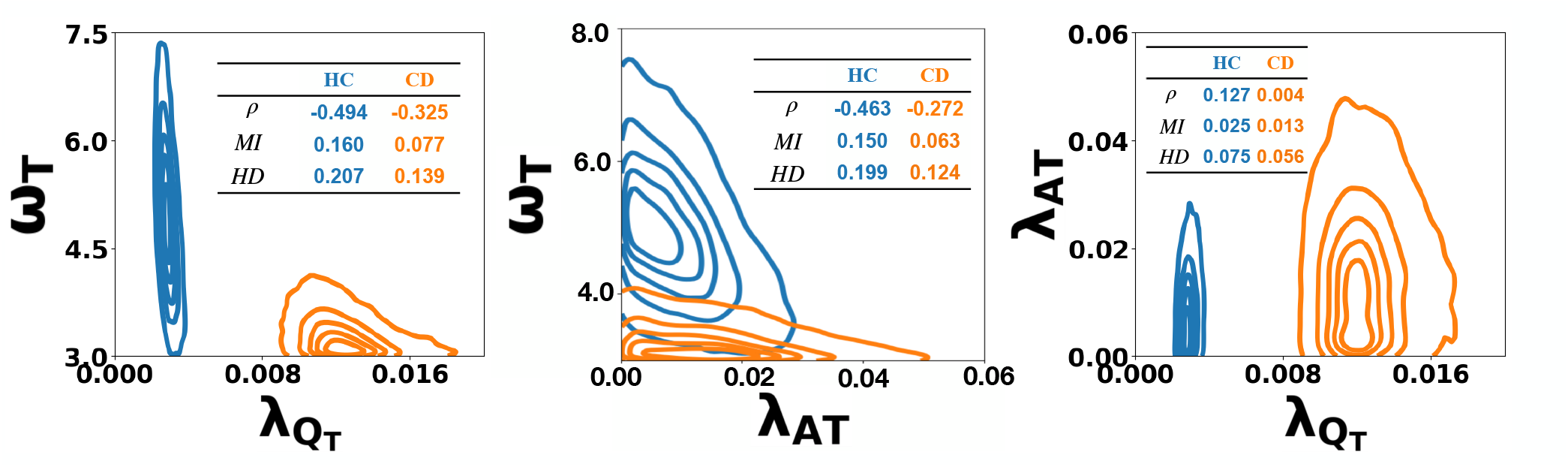
Joint posterior distributions reveal structured trade-offs among key task-level parameters. HC is plotted in blue. CD is plotted in orange. Inlet tables show estimated Spearman’s *ρ*, mutual information (MI), and Hellinger Distance (HD) values for HC vs. CD. for each parameter pair.

**Figure 7.**
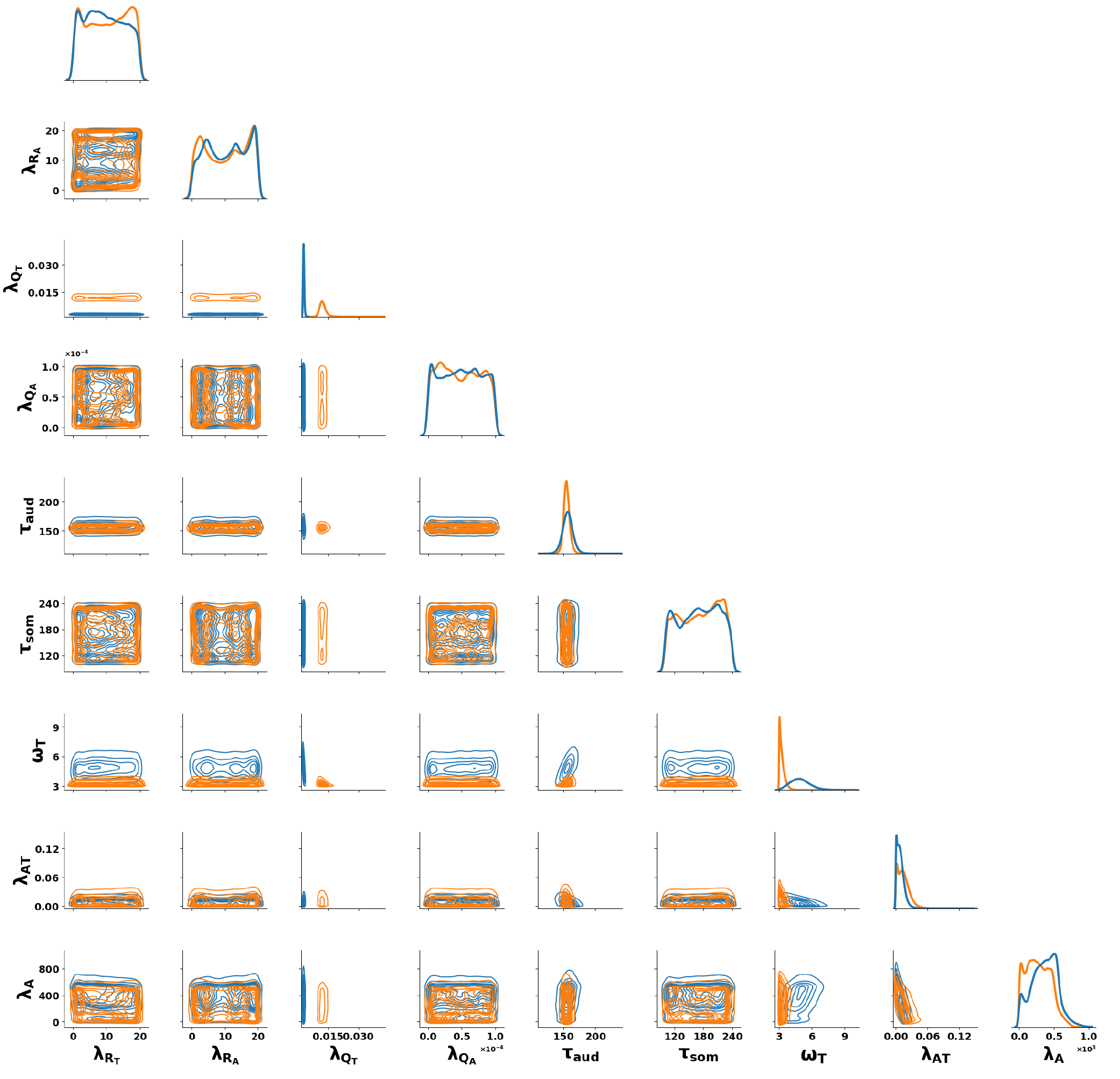
Joint posterior distributions over the parameters. HC in blue and CD in Orange.

A similar line of argument applies to the negative dependence between the timing of movement dynamics (*ω*_*T*_ ) and the multi-modal integration parameter (*λ*_*AT*_ ) (Figure 6; 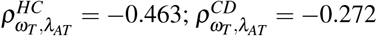). The negative dependence tells us that as the timing of movement dynamics gets slower (↓*ω*_*T*_ ), we can expect the multi-modal integration noise to get larger (↑*λ*_*AT*_ ). Indeed, the timing of movement dynamics value gets smaller (↓*ω*_*T*_ ; Glass Δ = −1.87) and the multi-modal integration noise value gets larger (↑*λ*_*AT*_ ; Glass Δ = 0.81) in CD. Taken together with the result that *λ*_*AT*_ did not contribute to significant behavioral differences in the single-parameter substitution anaylsis, this suggests that the contribution of *λ*_*AT*_ is relational rather than independently driving behavior. Elevated multimodal noise (*λ*_*AT*_ ) may help to accommodate the broader CD solution, but its isolated substitution does not produce a large enough trajectory change to rank as a primary behavioral driver. This interpretation is consistent with reports of slower timing of movement dynamics in CD [57], and impaired multimodal integration [21]. Again, although our work corroborates prior studies, those studies investigated the consequences of cerebellar damage through the lens of isolated mechanisms. Our work extends those accounts by providing a quantitative characterization – in terms of direction, mutual information, and strength of dependence – of their joint interaction.

Another result worth discussion is that across all parameter pairs, the CD mechanisms were less tightly coupled compared to HC (Figure 6; |*ρ*^*CD*^ | *<* |*ρ*^*HC*^|, *MI*^*CD*^ *< MI*^*HC*^, *HD*^*CD*^ *< HD*^*HC*^). This suggests that adults with CD may have aberrant speech compensation behavior not only because individual mechanisms have different values, but also because the mechanism interactions are less tightly coupled and may not be as ready to coordinate trade-offs. Although broader cerebellar motor-control literature shows that cerebellar ataxia involves increased variability and impaired coordination of motor synergies [59] and inter-joint dynamics [60], we could not find claims in the literature that show attenuated trade-offs with the specific mechanism pairs presented in this work. Thereby, this work may have generated a new hypothesis about CD having weakened coordination of the mechanisms forward modeling, timing, and multi-modal integration, which can be tested in future investigations.

Our results also revealed a trivariate structure organizing speech compensation control, with the timing of movement dynamics mechanism at the center of coordinating this subspace. The timing of movement dynamics mechanism (*ω*_*T*_ ) showed strong dependence, mutual information, and negative correlation with both the multi-modal integration parameter (*λ*_*AT*_ ) and the forward model parameter 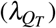 in both HC and CD. Whereas, 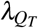 and *λ*_*AT*_ do not interact strongly with each other. This suggests that the timing of movement dynamics mechanism may serve as a central mechanism in the inferred regime, balancing trade-offs among multiple parameters in both HC and CD. This result is conceptually similar to the proposed coordinating role of the cerebellum [61], or to [24] where the authors propose that the cerebellum supports timing coordination between two pathways, albeit those pathways are different from the mechanisms highlighted in this work.

One question is whether the joint posterior analysis provides information beyond the marginal results, given that the marginal posteriors are themselves derived from the same joint posterior samples. If the joint analysis merely restated which parameters were larger or smaller in CD, then its interpretation would indeed be trivial. However, marginalization discards information about dependence: the same marginal distributions can be associated with positive, negative, independent, nonlinear, or funnel-shaped joint posterior geometries. To highlight alternative posterior interactions that could have explained the data, the joint posterior analyses could have revealed synergistic dependencies (i.e. a strong positive dependency: as 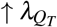 then ↑ *λ*_*AT*_ ) that could have explained the result that both of those mechanisms increased together. Another hypothetical result could have shown the forward model parameter 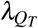 as a central mechanism instead of the timing of movement dynamics parameter *ω*_*T*_ . Other results could have revealed funnel-shaped posteriors, suggesting varying ranges of multi-parameter sensitivity. The joint analysis therefore does not simply reproduce the marginal group differences; it identifies the particular multivariate organization within which those differences occur. This organization does not establish a biological interaction, but it provides a more constrained model-based hypothesis about how the inferred mechanisms may compensate for one another.

In summary, our work suggests that disordered CD F1 compensation lies along a low-dimensional trade-off manifold involving timing of movement dynamics (*ω*_*T*_ ), forward model process noise scale 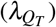, and multi-modal integration noise (*λ*_*AT*_ ) with CD differing not only in single mechanism values but also in multi-mechanism trade-off strengths and boundaries. Across analyses, the central result is that exaggerated compensation in cerebellar degeneration is explained by a noisier forward model 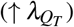, with slower timing of movement dynamics (↓ *ω*_*T*_ ) coordinating negative trade-off regimes with both the forward model 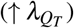 and multi-modal integration (↑ *λ*_*AT*_ ) mechanisms. Further, the strength between all of the multi-mechanism interaction pairs are weaker in CD compared to HC. This ability to recover joint structure across parameters allows investigators to disentangle contributions from parameters that even a priori have a similar effect on output behavior. This suggests that broader adoption of SBI-style joint inference could improve identifiability and interpretability in other neuroscience and behavioral studies that seek to quantify and disentangle neuroscience mechanisms.

### From isolated mechanisms to coordinated parameter regimes

These findings point to a broader limitation in the field. Computational functions are often investigated via a *ceteris paribus* approach—varying one candidate mechanism while holding all other mechanisms constant. Although this is valuable for generating interpretable causal predictions, most brain regions and their constituent subcircuits have been associated with multiple computations that may concurrently contribute to behavior. For example, prefrontal circuits contribute to working-memory maintenance, abstract-rule representation, and value coding [62, 63, 64]; hippocampal circuits contribute to spatial coding, temporal-sequence representation, and the construction of imagined experiences [65, 66, 67]; and basal-ganglia circuits contribute to action-value representation, habit formation, and movement vigor [68, 69, 70]. Investigating these functions individually can establish their possible contributions to behavior, but it cannot determine whether the underlying mechanisms act independently, compensate for one another, reinforce one another, or constrain the parameter ranges over which the other mechanisms can operate.

Systems neuroscience may therefore benefit from complementing single-mechanism perturbations with computational models that instantiate several candidate mechanisms simultaneously and estimate their joint contributions to behavior. Joint Bayesian inference, including simulation-based inference for complex and stochastic neural models, provides a means of estimating the full parameter space compatible with empirical observations rather than selecting a single best-fitting parameter set [71]. This approach can identify mechanistic trade-offs, synergistic dependencies, asymmetric constraints, and changes in coupling across populations, learning states, or task contexts. Such analyses complement, rather than replace, causal perturbation experiments, and can guide those experiments toward combinations of mechanisms that are jointly informative. Applied across cortical, subcortical, and cerebellar systems, this paradigm could move the field beyond cataloging functions associated with individual regions toward identifying the coordinated computational regimes through which neural systems generate behavior.

### Limitations

It is important to note limitations in our investigation. The first is that our hypotheses were evaluated within the context of a single speech motor task of compensatory responses to F1 perturbations during full trial vowel production. While F1 perturbation tasks are well-established probes of sensorimotor integration and correction [38, 72, 73], it does not encompass the full range of cerebellar contributions to speech. For example, if we were to model tasks with explicit delay, rhythm, or temporal sequencing, then parameters related to timing delays (e.g., *τ*_*aud*_, *τ*_*som*_) may prove more behaviorally relevant. Similarly, if our task perturbed speech articulators [74], then the parameters related to the articulatory modules (e.g. 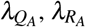, *λ*_*A*_) may have had more impact on the behavioral output. Thereby, our results speak to a subset of tasks, and should be interpreted within that scope.

### Future Directions

Although the present work highlights the cerebellum’s contributions to motor control, it is important to emphasize that motor control arises from distributed computations across multiple brain regions [1, 75, 76]. For example, regarding the task-state feedback control law, task representations are also associated with the orbitofrontal cortex [77, 78, 79], while speech-related feedback control laws are thought to involve the left ventral premotor cortex and primary motor cortex, supplemented by cerebellar projections [80, 81, 35]. The articulatory-state feedback control law is commonly attributed to the primary motor and sensory cortices [11]. Although the anatomical and functional literature, combined with our cerebellar degeneration cohort, motivated our emphasis on cerebellar mechanisms in this work, future studies may extend this approach to other brain regions and neurodegenerative disorders [82]. Future work may also integrate functional neuroimaging data (e.g., fMRI, MEG, DTI, SEM connectivity, etc.) to test whether individual differences in control parameters correlate with activity in cerebellar, sensorimotor, executive functioning, and auditory processing circuits [83, 84].

A future direction is to extend the present group-level framework to simulation-based inference at the individual-participant level. Group-level posteriors identify mechanisms that distinguish CD from healthy controls on average, but they may obscure heterogeneity within the CD population. In particular, some participants with CD show pronounced or atypical compensatory responses [38], whereas others exhibit responses that overlap with those of healthy adults [42]. Individual-level SBI could estimate a separate posterior distribution over mechanisms for each participant, allowing investigators to determine whether these behavioral differences arise from variation in forward modeling, movement dynamics, sensory integration, or distinct combinations of mechanisms. More broadly, moving toward individual-level inference may reveal mechanistic subtypes within CD and support more precise links among neural degeneration, computational dysfunction, and behavioral phenotype.

Another future direction is to investigate whether SBI can inform the design of model-guided brain stimulation therapies. If disordered speech compensation arises primarily from a shift in a single computational mechanism, stimulation protocols could be developed to modulate the neural circuitry supporting that mechanism. The present findings, however, suggest a more complex possibility: adults with cerebellar degeneration may differ not only in individual mechanism values, but also in how multiple mechanisms are coordinated within a broader control regime. Future studies could therefore test whether effective neuromodulation requires restoring coordination among forward modeling, multimodal processing, and the timing of movement dynamics, rather than normalizing any one computation in isolation. By combining individual-level SBI with neuroimaging and stimulation, investigators could identify parameter regimes associated with more adaptive speech behavior, map those regimes onto candidate neural targets, and evaluate whether stimulation shifts an individual’s inferred control system toward a more stable functional configuration. More broadly, this approach could establish computational models not only as explanatory tools, but also as treatment-planning frameworks for developing and personalizing stimulation protocols that act on interacting neural mechanisms.

Another extension would be to use SBI to evaluate the causal architecture of the computational model itself. Model components could be represented as nodes in a Bayesian belief network, while the strengths and temporal delays of connections between components are treated as parameters to be inferred from behavioral and neural data. The resulting posterior distributions could then assess whether the inferred dependencies and temporal ordering support the model’s assumed causal structure or favor alternative architectures. Although this approach would not establish causality from observational data alone, it could provide a principled means of comparing competing mechanistic models.

Together, these directions extend the present framework from group-level characterization of cerebellar mechanisms toward a broader, individualized, and translational account of motor control. Future studies can apply SBI across distributed neural systems, integrate neuroimaging to link inferred parameters to specific circuits, and perform individual-level inference to characterize mechanistic heterogeneity within CD. Combining these approaches with brain stimulation may ultimately enable computational models to guide personalized interventions that target not only altered mechanisms, but also their coordination within the broader motor-control system. In summary, future work may extend this framework to include more tasks, neurodegenerative groups, individual differences, causal experiments, mechanistic architectures, and multimodal neural recordings to further clarify how hidden processes in the human brain coordinate and drive behavior.

## Methods

We used an extended version of the Hierarchical Feedback-Aware Control of Tasks in Speech (FACTS) model to determine which neurocomputational mechanisms best explain speech responses in healthy adults and adults with cerebellar degeneration. The model represented nine parameters related to internal prediction, sensory-error processing, multimodal integration, feedback delays, feedforward control, and movement dynamics. We generated 1,002,080 simulations of first-formant (F1) responses to a +150-Hz auditory-feedback perturbation and used simulation-based inference to learn the relationship between model parameters and speech behavior. We then estimated separate parameter distributions for the observed healthy-control and cerebellar-degeneration trajectories, refining each estimate through four additional rounds of targeted simulations. We compared the groups’ inferred parameter distributions and evaluated the model using posterior predictive checks. Finally, we used parameter-substitution and marginal sensitivity analyses to measure how strongly each mechanism influenced the predicted speech response, and posterior-dependence analyses to identify coordinated relationships and trade-offs among mechanisms. A public version of the model and analysis pipeline is available at this GitHub repository

### Feedback Aware of Tasks in Speech: FACTS

The Hierarchical Feedback-Aware Control of Tasks in Speech (FACTS) Design C [34] was used, and extended, to generate simulated speech data for simulation-based inference. FACTS Design C implements a hierarchical state feedback control architecture that integrates task-level and articulatory-level representations, multi-modal sensory feedback, and internal predictions. The model includes tunable parameters that correspond to hypothesized cerebellar computations related to speech motor control including internal modeling, sensory error processing, multi-modal integration, delay processing, and timing of movement dynamics. The FACTS model was extended to include delays in the state estimators *τ*_*aud*_ and *τ*_*som*_ separate from sensory feedback delays 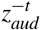 and 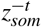. The FACTS model was also extended to include noise scales in the articulatory state feedback control law *λ*_*A*_ and the artic-to-task transformation *λ*_*AT*_ . The FACTS model implementation, parameter priors, simulation scripts, empirical datasets, and plotting routines are available in Extended Data 1.

### FACTS Parameters: Uniform Prior Boundaries

We defined a 9-dimensional parameter space corresponding to key cerebellar computations (Table 1) including: internal forward models 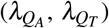, feedforward control (*λ*_*A*_), error detection via auditory and somatosensory noise scaling 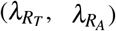, multi-modal integration via the articulatory-to-task state noise scale (*λ*_*AT*_ ), timing delays via the articulatory- and task-level delays (*τ*_*som*_, *τ*_*aud*_), and timing of movement dynamics (*ω*_*T*_ ). Each parameter *θ*_*j*_ was assigned a uniform prior range *θ*_*j*_ ∼ *Uni f orm*(*a*_*j*_, *b* _*j*_) (Table 2).

**Table 2.** Prior boundaries for model parameters.

| Parameter | Prior Min | Prior Max |
| --- | --- | --- |
| $\lambda_{R_T}$: Task State Estimator Auditory Noise Scale | 1e-8 | 2e1 |
| $\lambda_{R_A}$: Articulatory State Estimator Somatosensory Noise Scale | 1e-8 | 2e1 |
| $\lambda_{Q_T}$: Task State Estimator Process Noise Scale | 1e-8 | 5e-2 |
| $\lambda_{Q_A}$: Articulatory State Estimator Process Noise Scale | 1e-8 | 1e-4 |
| $\tau_{aud}$: Task State Estimator Auditory Delay | 1e2 | 2.4e2 |
| $\tau_{som}$: Articulatory State Estimator Somatosensory Delay | 1e2 | 2.4e2 |
| $\omega_T$: Natural Frequency of the Task State Feedback Control Law | 3 | 1e1 |
| $\lambda_{AT}$: Articulatory-to-Task State Noise Scale | 0 | 2e-1 |
| $\lambda_A$: Articulatory State Feedback Control Law Noise Scale | 0 | 1e3 |

**Table 3.** SBI training parameters.

| Parameter | Value |
| --- | --- |
| Training Batch Size | 32 |
| Force First Round | True |
| Discard Prior Samples | True |
| Density Estimator | 'nsf' |
| Learning Rate | 5e-6 |

**Table 4.**
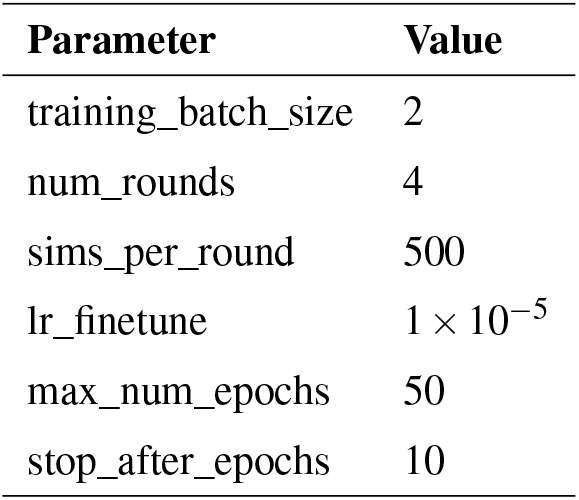
Shared fine-tuning hyperparameters used for multi-round SBI in both conditions.

### Prior Sampling and FACTS Simulations

1,002,080 parameter vectors ***θ*** _***i***_ were generated, where each element in the vector was sampled from the uniform prior distribution *θ*_*j*_ ∼ *Uni f orm*(*a*_*j*_, *b* _*j*_). Each parameter vector was used as input to generate corresponding *F*1_*i*_ speech formant trajectories. Parameter and F1 pairs 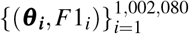 were created and used to train the SBI neural conditional density estimator.

FACTS created F1 trajectories at 5ms resolution such that 100 frames equals 500ms. To mirror the empirical data, whole trial perturbations and 500ms trial durations were simulated. To ensure that the simulation outputs were stable when the perturbation was applied, we simulated 700ms trials and applied the perturbation at 200ms from onset; after all simulations were generated we then created a copy of the dataset, removing the first 200ms of the trials to thereby simulate 500ms whole trial perturbations. Whole trial F1 auditory perturbations of +150 Hz were applied to mirror the empirical data experimental design [38].

### Empirical data

The behavioral data from Parrell et al., 2017 [38] were used, which show HC and CD speech compensations to random whole trial F1 perturbations of *±*150Hz. Trajectories were averaged over nineteen CD participants and fourteen HC participants, with experimental blocks containing 170 trials. Auditory feedback was altered using the Feedback Utility for Speech Production (FUSP) system [85]. For our experiment, to make RMSE values comparable between empirical HC and CD trajectories vs. SBI-derived HC and CD trajectories, 1Hz was subtracted from the empirical CD F1 mean trajectory so that the CD and HC pre-compensation frames were more closely overlapping. This alteration is justified because the CD and HC F1 trajectories were not significantly different before compensation onset, and modeling initial F1 was outside of the scope of this study. This decision obviates pre-compensation F1 speech segments from erroneously inflating RMSE for one class over the other, and thereby make F1 RMSE values more readily comparable. For more information on the empirical data, please refer to [38]

### SBI Training and Inference

Simulation-Based Inference (SBI) [46], version 0.22.0, was used to infer control parameter values underlying speech motor behavior. Using the simulation pairs 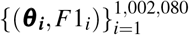, a neural conditional density *q*_*φ*_ (*θ* | *F*1) was trained. Given empirically observed data 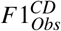 and 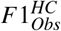 from [38], parameter posterior distributions were approximated 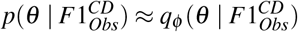 and 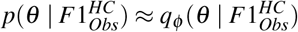. The Sequential Neural Posterior Estimation (SNPE) inference algorithm was used. During SBI training the following parameters were set:

The model converged after 217 epochs with a best validation performance of -2.7097 nats. 100,000 samples were drawn from the posterior, and the modes of the marginal parameter posterior distributions were calculated and stored 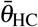 and 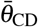. The modes were used to for posterior predictive checks, and subsequent analyses such as parameter posterior analysis, single-parameter substitution analaysis, and marginal sensitivity analysis.

### Multi-round SBI

Because the initial single-round SBI analysis produced posterior parameter estimates whose simulated F1 trajectories fell outside the standard error of the mean (SEM) bounds reported in Parrell et al., we performed a multi-round SBI refinement procedure to improve the correspondence between simulated and empirical behavior. In this approach, the posterior inferred in one round was used to guide simulation and posterior refinement in the next round, thereby concentrating samples in regions of parameter space that were more consistent with the observed data.

Multi-round inference was performed separately for the healthy control (HC) and cerebellar degeneration (CD) conditions using the same fine-tuning hyperparameters for both groups, shown in the table below.

### Parameter Posterior Distribution Analysis

Given formants 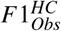 and 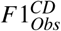, 100,000 parameter vectors were sampled from the estimated posterior distribution to build an estimated sample distribution 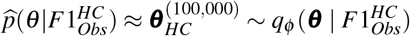 and 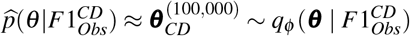. Marginal posterior distributions were calculated for each parameter *θ*_*i*_, indexed at i, and the modes of the marginal posteriors were reported as the maximizer of the estimated marginal posteriors:

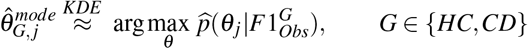

where KDE denotes the Kernal Density Estimator of the Scipy package [86]. Violin plots and joint posterior PairGrid plots were made using the seaborn package [87] in python. Glass’s delta effect sizes Δ_Glass_ were calculated for each parameter.

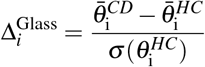

where 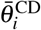 and 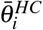 is the mode for parameter indexed at i for each group, and 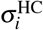 is the standard deviation of parameter indexed at parameter i of the Healthy Control group. Posterior uncertainty was summarized using the highest-density interval (HDI) at probability mass *h* (here *h* = 0.95), defined as the minimum-width credible interval:

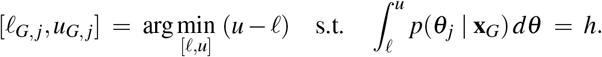

HDIs were computed from 100,000 posterior samples using `arviz.hdi` (ArviZ) [88, 89].

Non-parameteric two-sided Mann-Whitney p-values were calculated to compare HC vs. CD of each marginal posterior distribution. Given the large samples size N=100,000 per group, p-values can become extremely small, and values below numerical precision were reported as *<* 10*e*^−300^.

### Single-parameter substitution analysis

To quantify the contribution of each parameter to CD speech behavior, we performed a single parameter substitution analysis. The analysis asks, for each parameter indexed at *i*, how much the CD F1 trajectory changes when only that parameter is replaced by its healthy control (HC) counterpart, while all other parameters remain at their CD values. This isolates each parameter’s marginal influence on CD behavior in the neighborhood of the inferred CD setting.

For both the HC and CD groups, we computed posterior parameter vectors 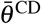 and 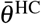 containing the respective groups’ marginal posterior modes. We then defined single-parameter substituted vectors 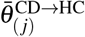 by replacing one CD parameter mode with its corresponding HC mode at index j, yielding:

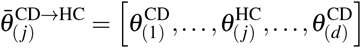

and *d* = 9 for the total number of parameters.

A mean 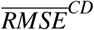 value was calculated by first comparing simulated trajectories 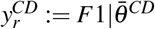 to the target trajectories 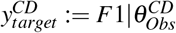 [38], and then taking the mean of each trajectory’s 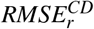 across the total number of simulation runs R=25 indexed by r.

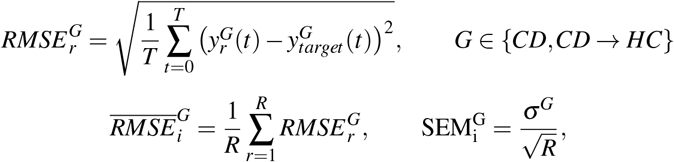

To quantify the magnitude of change induced by substituting parameter *i*, we computed Glass’s Δ using the CD baseline as the control distribution:

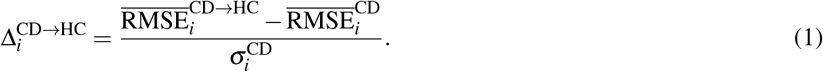

Here, 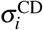 is the sample standard deviation of 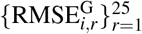. Positive values of 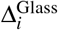 indicate that replacing 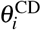 with 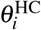 increased error relative to the CD baseline, whereas negative values indicate decreased error.

### Marginal Sensitivity Analysis

This procedure systematically deviates each parameter around its posterior mode while holding all other parameters constant, enabling us to evaluate how changing a parameter’s value, with respect to its central tendency and variance, affects model output. Starting with the parameter posterior modes for each group, 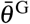 for *G* ∈ *{HC,CD}*, we estimated a per-parameter posterior standard deviation from posterior draws. 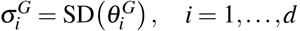. We then used standard deviation coefficients *i*: *p* ∈ *{*−3, −1.5, 0, 1.5, 3*}* while holding all other parameters fixed, yielding:

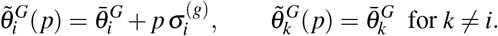

So only the parameter of interest, indexed at i, was changed, whereas the rest of the parameters, indexed at k and *k* ≠ *i*, were kept to their original mode values. Because several FACTS parameters are constrained to be non-negative in our implementation, any condition producing 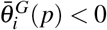 for either group was treated as infeasible and excluded from analysis (plotted as empty barplot columns or missing F1 trajectory values).

Each 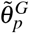 was passed through the FACTS model 25 times to generate 25 simulated F1 trajectories for visualization and statistical analysis. For each parameter, we visualized sensitivity using two complementary summaries. First, we plotted the mean fit error, 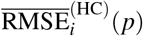 and 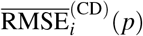, as a function of the perturbation level *p*, using side-by-side bar plots for the HC and CD groups with error bars given by the standard error of the mean, 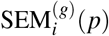. This representation provides a local measure of how strongly deviations in parameter *i* affect overall agreement between simulated and observed trajectories. Second, to characterize the time-resolved consequences of each perturbation, we plotted the mean simulated trajectory:

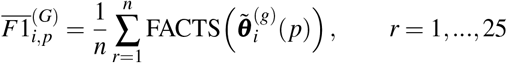

and overlaid the corresponding observed trajectory 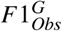. This visualization identifies when, and in what direction, perturbing parameter *i* induces systematic deviations in the predicted F1 dynamics. Together, these visualizations yield an interpretable, local characterization of parameter sensitivity in each group, enabling identification of parameters to which the modeled F1 trajectories (and associated RMSE fit) are most responsive.

By plotting RMSE as a function of deviation magnitude for each parameter, we assessed the local behavioral sensitivity to parameter variation. Parameters with a steep RMSE gradient were interpreted as exerting strong influence over the model’s output dynamics. This analysis complements the single-parameter substitution analysis by revealing how the model’s behavior changes in the local neighborhood of the inferred parameters, rather than a single-parameter cross-group substitution.

### Posterior dependence analysis

To quantify pairwise dependence among the inferred parameters, we analyzed posterior samples for each parameter pair (*θ*_*i*_, *θ*_*j*_) separately within each group. For a given pair, we assessed three complementary quantities: Spearman’s rank correlation coefficient, mutual information, and Hellinger distance between the pairwise joint posterior and the product of its marginal posteriors.

#### Posterior samples

For each group, posterior samples were drawn from the inferred posterior distribution conditioned on the corresponding observed summary statistic. For each parameter pair (*θ*_*i*_, *θ*_*j*_), these samples were used to estimate both the pairwise joint posterior *p*(*θ*_*i*_, *θ*_*j*_ | *x*) and the marginal posteriors *p*(*θ*_*i*_ | *x*) and *p*(*θ*_*j*_ | *x*).

#### Spearman’s rank correlation

We first quantified monotonic dependence between *θ*_*i*_ and *θ*_*j*_ using Spearman’s rank correlation coefficient, denoted *ρ*. Given posterior samples

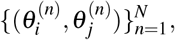

we replaced each variable by its rank and computed the Pearson correlation of the ranked samples. Spearman’s *ρ* ranges from −1 to 1, with negative values indicating that larger values of one parameter tend to co-occur with smaller values of the other, positive values indicating the opposite, and values near zero indicating weak monotonic dependence. We report Spearman’s *ρ* as a descriptive measure of the orientation and strength of posterior trade-off structure.

**Kernel density estimation of joint and marginal posteriors**

To compare the pairwise joint posterior against the factorized approximation implied by posterior independence, we estimated the joint density *p*(*θ*_*i*_, *θ*_*j*_ |*x*) using a two-dimensional Gaussian kernel density estimator (KDE) fitted to posterior samples. We used the scipy stats KDE package:

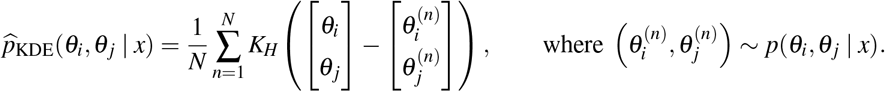

where *K*_*H*_ denotes a Gaussian kernel with bandwidth matrix *H*. The quantity in parentheses represents the displacement between the grid evaluation point (*θ*_*i*_, *θ*_*j*_) and the *n*th posterior sample 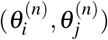, which determines that sample’s contribution to the estimated density. Intuitively, instead of treating each sample as an infinitely sharp point, the KDE places a smooth hill around it. Adding all of these hills produces a smooth surface. High regions of the surface indicate parameter combinations supported by many nearby posterior samples, while low regions indicate combinations with little posterior support. All densities were evaluated on a regular grid and numerically normalized so that they integrated to 1:

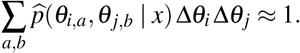

The implementation is available in the Github link.

#### Mutual information

We quantified non-factorization of the pairwise joint posterior using mutual information,

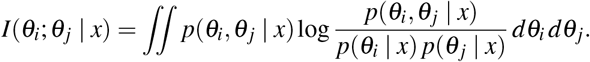

Mutual information is equal to zero if and only if the joint posterior factorizes as

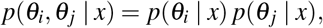

and is positive when the two parameters are posteriorly dependent. In practice, mutual information was approximated numerically on the KDE evaluation grid:

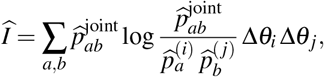

where 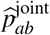 denotes the estimated joint density at grid cell (*a, b*), and 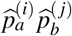 denotes the product of the estimated marginals at that cell. Mutual information was reported in nats.

#### Hellinger distance

As a complementary measure of discrepancy between the joint posterior and the product of marginals, we computed the Hellinger distance,

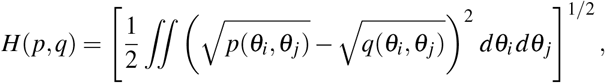

with

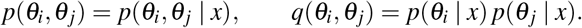

The Hellinger distance takes values in [0, 1], with *H* = 0 indicating exact equality between the joint posterior and the factorized approximation, and larger values indicating greater discrepancy. Numerically, this quantity was approximated on the same grid used for the KDE-based density estimates:

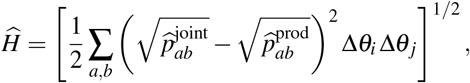

where 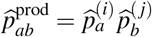.

#### Interpretation

Together, these three quantities summarize complementary aspects of posterior dependence. Spearman’s *ρ* captures the direction and monotonic strength of pairwise dependence, mutual information quantifies the extent to which the joint posterior fails to factorize, and Hellinger distance measures the overall discrepancy between the joint posterior and the corresponding product of marginals. Larger absolute values of *ρ*, larger mutual information, and larger Hellinger distance all indicate stronger pairwise posterior dependence.

## Acknowledgements

This work was funded by awards from the National Institutes of Health (P50DC019900, R01NS100440, R01DC017091, and R01DC017696 to S.S.N. and J.F.H.) and the National Science Foundation (2034836 to A.L.P.). A special thanks to Rich Ivry and Pascal Perrier for insightful comments.

## Author contributions statement

A.P., K.K., B.P, J.G., V.K., R.I., S.N., J.H conceived the experiment(s),; A.P., K.K., N.C. conducted the experiment(s), A.P., K.K., J.G.; A.P. K.K., B.P., J.G., V.R, R.R, N.C., K.B., S.N., J.H. analyzed the results.; A.P., K.K., B.P., J.G., V.R, R.R, R.I., N.C., K.B., S.N., J.H. wrote and reviewed the manuscript.

## Additional information

To include, in this order: **Accession codes** (where applicable); We declare that none of the authors have competing financial or non-financial interests as defined by Nature Portfolio.

The corresponding author is responsible for submitting a competing interests statement on behalf of all authors of the paper. This statement must be included in the submitted article file.

## Supplemental

